# Fun-Sized Library Prep: Miniaturization is a valid method for per-sample cost reduction in targeted sequencing of angiosperm DNA

**DOI:** 10.1101/2025.09.17.676862

**Authors:** Madison R. Bullock, Mohamed Fokar, Rylee D. Creek, Eliot A. Stevens, Matthew G. Johnson

## Abstract

**Premise:** Genomic analysis of population structure is important to the conservation of plant species of concern. A limitation of using genetic information in conservation is the cost of obtaining large datasets. Targeted sequencing and low-volume robotic liquid handlers can reduce library preparation reaction volumes and costs.

**Methods:** We used targeted sequencing via Angiosperms353 to obtain data for 768 samples, 18 of which were identical, at 0.5X and 0.1X reaction volumes. We calculated quality and quantity control statistics to compare the effects of tissue age and library reaction volume on sequencing results for on-target nuclear and off-target plastid genes.

**Results:** Library miniaturization to 0.1X reduces costs and generally performs comparably to 0.5X libraries. In the full dataset, Tenth-Volume Only libraries showed smaller insert sizes and fewer genes with mapped sequences, but no reduction in mapped reads. In the Overlap Set, 0.1X had equal or improved performance with no significant decrease in sequencing efficiency. Differences by tissue type likely reflected sampling variation.

**Conclusions:** Miniaturization to 0.1X substantially reduces per-sample costs while maintaining comparable sequencing quality across fresh and herbarium angiosperm DNA. Overall, miniaturization provides a reliable, cost-effective approach for targeted sequencing, increasing the feasibility of using herbarium collections and enabling broader access to population-level genomic studies.

## INTRODUCTION

Conservation genetics is essential to maintaining important genetic diversity within populations. In the short term, genetic diversity affects the fitness of the population via increased performance, decreased inbreeding depression, and decreased susceptibility to pathogens (Smithson and Lenne, 1996; Hughes et al., 2008; Wan et al., 2022). In the long term, the maintenance of genetic diversity in plant taxa preserves evolutionary potential essential to species survival through the maintenance of biodiversity and subsequent ability to change in response to the environment (Frankham, 2005; Frankham, 2010). However, there is a gap between conservation genetics research and its application in management plans, leading to underutilization of genetic information in conservation decisions (Magris et al., 2016; Cook and Sgrò, 2017; Rose et al., 2018). Studies have found that this gap is largely due to lack of funds available for conservation genetics research as these studies typically require large, multi-individual datasets (Bradshaw and Brook, 2010; Taylor et al., 2017; Taft et al., 2020). Therefore, despite recent declines in high-throughput sequencing costs, there is a need for further decreasing per-sample costs to increase accessibility of genetic research to conservation management.

Conventionally, genetic data can be generated by a variety of methods such as whole genome, restriction digest sequencing, and target capture sequencing (reviewed in McKain et al., 2018). But because of expensive field work trips to collect samples, the price of reagents for full volume (1.0X) library preparation reactions, and the price of sequencing itself, it can be costly to generate datasets of the size needed for robust inference (Hale et al., 2020). Target capture sequencing was developed to selectively obtain orthologous regions of interest across multiple individual samples in a more cost-effective manner compared to other sequencing methods (Olson, 2007) – on average, up to three times cheaper than whole genome sequencing (Bewicke-Copley et al., 2019). In plants, target capture typically enriches libraries using probes designed from the exon regions of genes across a group of interest, which can range from single genera (*Artocarpus*; Gardner et al., 2015) to families (Euphorbiaceae; Villaverde et al., 2018), or orders (Bryopsida, Liu, Johnson, et al., 2019). An additional advantage of targeted sequencing approaches is the possibility of sequencing herbarium specimens (Brewer et al., 2020) despite degraded DNA to compare across spatiotemporal scales (Ding et al., 2020; Hagen et al., 2021).

Several “universal” probe sets – such as Angiosperms353 (Johnson et al., 2019) and Compositae1061 (Mandel et al., 2014) are available for reduced cost. In 2020, Hale et al. described a series of measures aimed at cutting per-sample costs when pooling samples for the Illumina MiSeq, with an estimated $30 USD per sample for consumables and reagents. Applications of this reduced-cost approach have extended beyond phylogenetic studies to use targeted exons and flanking non-coding regions to estimate demographic parameters (e.g. Slimp et al., 2021) or study the association between genetic and geographic variation within species (e.g., Beck et al., 2021). However, extending these advances to conservation genetics projects would require hundreds of samples, meaning even the cost reductions described by Hale et al. (2020) are insufficient to make within-species questions feasible.

Two stages of the target enrichment sequencing workflow that carry most of the per-sample costs are library preparation and sequencing. We propose to address both through massive multiplexing on the NovaSeq 6000 platform (Illumina, San Diego, California, USA) and library miniaturization. Currently, there are five sets of 96 NEBNext Multiplex Oligos for Illumina Unique Dual Index Primer Pairs (New England Biolabs, Ipswich, Massachusetts, USA). During library preparation, these primer pairs are ligated to the DNA as a barcode that can be used to demultiplex sequence data generated from the two flow-cell lanes of the NovaSeq 6000 platform. So, a total of 960 samples (480 per lane) can be sequenced on the same sequencing run, generating up to ten times the data of a standard Illumina MiSeq run.

Furthermore, library preparation miniaturization can address the need for decreased per-sample costs via lower requirements for consumables and reagents. Employing automated liquid handling robots, library miniaturization scales down the volume of reagents in a library preparation reaction. Advantages include processing speed, pipette accuracy, and maximization of reagent usage (SPT Labtech, 2023). Several studies have successfully used library preparation miniaturization of single-cells (e.g., Mora-Castilla et al., 2016; Johnson et al., 2022) and viruses (e.g., Yángüez et al., 2020; Karthikeyan et al., 2022; Pillay et al., 2023) prior to sequencing.

However, there have been fewer applications using plant tissues. For example, Haveman et al., 2022 used 0.1X library miniaturization prior to RNA sequencing for analysis of plant growth response in KSC Fixation Tubes to suborbital spaceflight. Hamala et al., 2023 also miniaturized their Cochlearia DNA libraries for long-read sequencing and identification of genomic structural variants related to polyploidy. Nevertheless, no studies have used library miniaturization on herbarium tissue-derived DNA or for targeted sequencing in plants to our knowledge.

We aimed to determine whether: 1) library miniaturization affects post-target capture sequencing data quality and quantity of angiosperm DNA, 2) library miniaturization works on herbarium tissue as well as it does on fresh plant tissue, and 3) library miniaturization for target capture sequencing is a valid method for costs reduction. Therefore, the purposes of this study were to determine the effects of library miniaturization on post-sequencing data quality and quantity, to analyze differences in library miniaturization of fresh and herbarium tissues, and to calculate the cost-savings of a 0.1X and 0.X approaches in comparison 1.0X approach. We hypothesized that library miniaturization does not affect post-sequencing data quality or quantity, and we tested this hypothesis via two analyses: 1) a comparison of post-sequencing statistics for two full target capture sequencing runs of containing herbarium or fresh tissue from 768 samples, and 2) a direct comparison of 18 libraries sequenced at both 0.5X and 0.1X reaction volumes. We also determined the overall cost of each method as a percentage of full volume (1.0X) costs.

## METHODS

### Sampling and DNA Extraction

For this study, we utilized an opportunistic set of angiosperm specimens being sampled for various projects including but not limited to ethnobotany, phylogeography, conservation genomics, population genomics, taxonomic resolution, and global change research. This diverse sample set included 1,518 individual plant specimens across 243 species, 176 genera, and 63 families ranging in age from less than a year to over 110 years (Appendix 1). Though some samples were from fresh tissue, we also sampled from herbarium specimens. Overall, the samples can be classified in three categories: Half-Volume Only (using methods from Hale et al., 2020), Tenth-Volume Only (using miniaturization methods described below), and the Overlap Set of 18 samples sequenced using both methods.

We sampled 1 cm^2^ from each individual and placed the tissue in 1.1mL strip tubes arranged in sixteen 96-tube plates with two steel ball bearings and capped them. The samples were then flash-frozen using liquid nitrogen in a SPEX Cryo-Station (SPEX SamplePrep, Metuchen, New Jersey, USA) and ground into powder by shaking each plate at 1500RPM for one minute in the SPEX Geno/Grinder MiniG tissue homogenizer. This process was repeated up to three times to ensure complete pulverization of the tissue. We extracted DNA through a CTAB/chloroform method (Doyle and Doyle, 1987) with a few modifications found to maximize the yield of DNA from older tissue such as from herbarium specimens (Brewer et al., 2019).

These include incubation of samples in the CTAB buffer for 24 hours instead of one hour, and precipitation of DNA in isopropanol at -20°C for a week instead of 24 hours. For quality control, we quantified the resulting DNA using a Qubit Fluorometer (Thermo Fisher Scientific, Waltham, Massachusetts, USA) and determined average fragment size per sample through gel electrophoresis. Finally, we performed an initial cleanup of the DNA using a 1.1X volume of homebrew SPRI beads (Rohland and Reich, 2012) to remove impurities and potential secondary metabolites.

### Library Preparation, Hybridization, and Sequencing

We prepared dual-indexed Illumina sequencing libraries from the extracted DNA via the NEBNext Ultra II DNA kit (New England Biolabs, Ipswich, Massachusetts, USA) following the manufacturer’s protocol with two modifications. For Half-Volume Only samples, we manually prepared 768 libraries at 0.5X reaction volumes using the protocols described in Hale et al. (2020). Depending on quantification, most samples were either diluted or concentrated to achieve as close to a per sample quantification of 200ng as possible prior to library preparation. The Tenth-Volume Only libraries we prepared for the second sequencing run were at 0.1X reaction volumes made possible by SPT Labtech Dragonfly and Mosquito (SPT Labtech, Hertfordshire, UK), two low-volume automated liquid handling robots. These samples were not strictly normalized as such small sample quantities were required by the protocol and preliminary tests of the protocol found normalization led to less effective library preparation when compared to no normalization (data not shown). Furthermore, we employed a few modifications found to increase library quantity and quality at low reaction volumes: adaptor diluted 1:10 in 10mM NaCl, 500μL of adaptor, 500μL of USER enzyme, 0.8X AMPure XP bead (Beckman Coulter, Brea, California, USA) concentrations during both cleanup steps, and eight PCR cycles. For direct comparison, an Overlap Set of 18 DNA extractions were prepared into libraries at both reaction volumes. Across all protocols, samples that were primarily comprised of fragments larger than 500bp as quantified on an agarose gel were enzymatically sheared via fragmentase as part of end repair. We quantified concentrations of the Half-Volume Only libraries using a Qubit Fluorometer. However, due to the ability of the liquid handling robot to transfer nanoliter amounts, we quantified the Tenth-Volume Only libraries using the Invitrogen Quant-iT high sensitivity kit (Invitrogen, Waltham, Massachusetts, USA) and a Biotek plate reader (Biotek, Winooski, Vermont, USA). We also determined fragment size distribution for a subset of 2-3samples per library preparation plate that covered the entire of the library concentration range from each reaction volume category using the Agilent TapeStation 2200 (Agilent Technologies, Santa Clara, California, USA).

Following a secondary quality control step, we pooled libraries based on DNA concentration at a maximum of 24 samples per pool for Arbor Biosciences’ Angiosperms353 myBaits hybridization capture (Arbor Biosciences, Ann Arbor, Michigan, USA). This resulted in 32 pools of libraries prepared at both 0.5X and 0.1X volumes. We then concentrated the pools to the Angiosperms353 input volume of 7μL via centrifuging in a Thermo Fisher Savant DNA 120 SpeedVac Concentrator for 5 to 45 minutes, dependent on initial volume. We then completed the target capture reactions following the manufacturer protocol with two -modifications: dilution of the RNA probes (1:3), as suggested in Hale et al. 2020, and 14 cycles of PCR enrichment following hybridization, which is the manufacturer-recommended maximum number of cycles. Once again, we quantified each pool via Qubit fluorometer and determined fragment size distribution using the Agilent TapeStation 2200. Subsequently, the pools were combined once again by total DNA concentration of each pool into two pools per reaction volume category before sequencing on the Illumina NovaSeq 6000 platform (Illumina, San Diego, California, USA).

### Cost Comparison of Library Reaction Volumes

We determined if reduction of library reaction volumes to 0.1X notably decrease financial and temporal costs of library preparation at three reaction volumes: 1.0X (full volume), 0.5X (half volume), and 0.1X (tenth volume). This includes all consumables (e.g., pipette tips, plates, tubes), library preparation reagents, quality control reagents, and total hands-on time needed to make a library from a DNA sample. Note that the costs are specific to Texas Tech University, the institution where this study was conducted, and may be subject to change dependent on the vendor.

### Data Processing and Statistical Analyses

We first processed the resulting sequence data via fastp (Chen et al., 2018) to remove the poly-G tail leftover from sequencing on the Illumina NovaSeq platform. This program also allowed us to estimate average insert fragment size via its JSON output files, with the limitation that fastp can only estimate insert sizes for fragments that are less than twice the read length.

Then, we assembled Angiosperms353 genes from the cleaned reads via HybPiper, a program for assembling genes, extracting both intronic and exonic sequences, and identifying paralogs (Johnson et al., 2016). Angiosperms353 genes include coding sequences of the targeted 353 genes as well as “supercontigs” containing exons and their flanking non-coding regions. Using HybPiper (Johnson et al., 2016), we generated the summary file of recovery statistics and a heatmap of per-sample percent gene recovery for each sequencing run and library reaction volume. We were specifically interested in the impact of library reaction volume on the insert fragment size, number of reads mapped, number of genes recovered by HybPiper, and total number of base pairs recovered for both exons only and “supercontigs.” The original description of the HybSeq technique (Weitemeier et al., 2014) describes simultaneous recovery of targeted nuclear regions and organellar loci from the “off target” regions. To test the impact of miniaturization and the NovaSeq multiplexing on plastid gene recovery, we used the angiosperm-wide target file from Pokorny et al. (2024) (available at https://github.com/mossmatters/plastidTargets) to recover plastid loci from the Overlap Set sequences.

To test for the direct consequence of library reaction volume on sequence data generation, we compared the average insert fragment size, reads mapped, and genes with sequences of 768 Half-Volume Only samples to 768 Tenth-Volume Only samples via a Welch Two Sample T-test from the stats package in R (R Core Team, 2021). Based on prior knowledge of reduced insert fragment sizes in Tenth-Volume Only libraries gathered during pre-sequencing quality control, we used fastp (Chen et al., 2018) and python (Van Rossum and Drake, 1995) to extract the average insert fragment size from the HybPiper-generated (Johnson et al., 2016) FASTA files of 768 Half-Volume Only and 768 Tenth-Volume libraries. To do a more direct comparison of the listed metrics as well as total number of base pairs recovered for “supercontigs,” we tested differences in the Overlap Set of 18 samples that were sequenced at both 0.5X and 0.1X library reaction volumes via a Welch’s Two Sample T-test. Finally, to assess whether any significant differences could be contributed to the tissue type the samples were derived from (herbarium or fresh) at each library preparation reaction volume, we utilized a one-way ANOVA followed by post-hoc analysis with a Tukey Honestly Significant Difference test, a two-way ANOVA to determine individual effects of tissue age and library volume, as well as their interaction, both with the stats package (R Core Team, 2021) and a power analysis of a subset sample of 100 libraries from each of the four groups.

## RESULTS

Within our complete data set of 768 Half-Volume Only and 768 Tenth-Volume Only libraries, we determined that library miniaturization to 0.1X reaction volumes significantly reduces average insert fragment size in number of base pairs comparatively to 0.5X reaction volumes (Figure 1A; T-test; t_1196.6_ = -12.402; P < 0.001). There was no significant difference in reads mapped (Figure 1B; T-test; t_1247.4_ = 1.556; P = 0.12), and a significant decrease in genes with mapped sequences when volumes were decreased to 0.1X (Figure 1C; T-test; t_1464.4_ = -6.82; P < 0.001).

**Figure 1.**
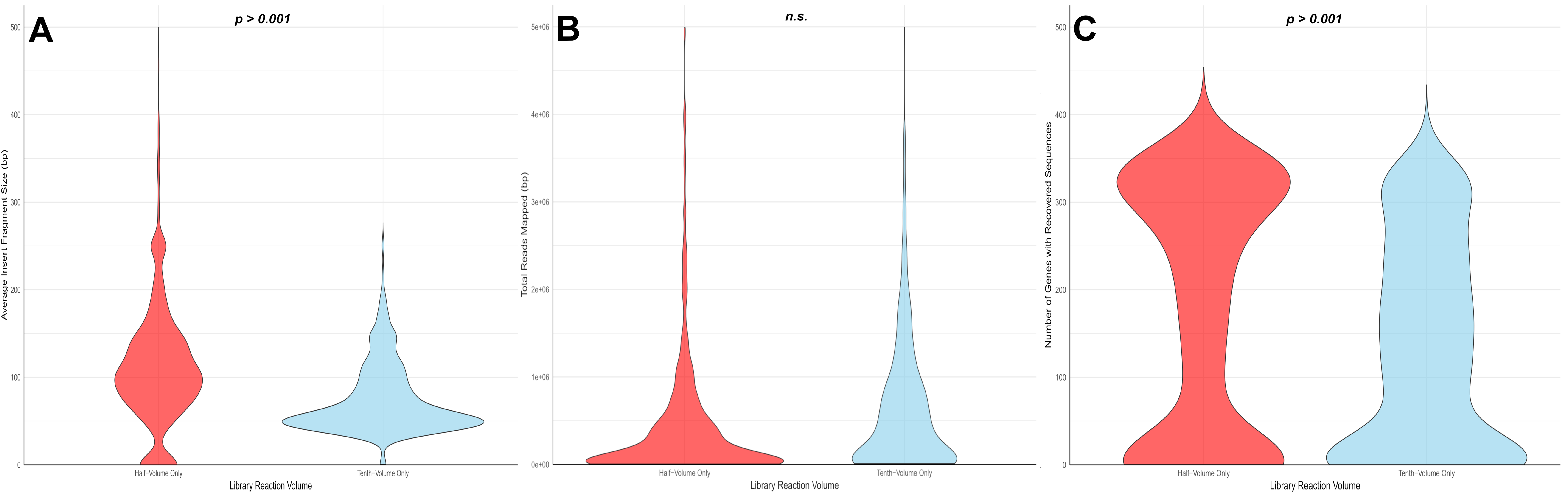
Average insert fragment sizes and genes with sequences are significantly lower in the Tenth-Volume Only libraries (blue) compared to the Half-Volume Only libraries (red). However, there is no significant difference in total reads mapped between Half-Volume Only and Tenth-Volume Only libraries. A. T-test; tl 196.6 == -12.402; P < 0.001. B. T-test; t1247.4 == 1.556; P == 0.12. C: T-test; t1464.4 == -6.82; P < 0.001.

To determine the direct effects of miniaturizing library preparation reactions on the generation of angiosperm targeted sequences, we chose 18 samples of DNA we had previously sequenced with 0.5X reaction volumes, and resequenced them using 0.1X reaction volumes as our Overlap Set. We then created heatmaps of the percent per gene recovered across all 353 targeted genes and across the 18 Overlap Set samples (Figure 2). Based on a visual comparison, we see that there is some variation on the individual sample scale as some samples that fail in one method perform well in the other. But there is no overall clear pattern based on reaction volume or library miniaturization. Within our Overlap Set, and contrary to the complete, opportunistic data, we found that library miniaturization to 0.1X had a significant positive effect on total reads mapped (Figure 3B; T-test; t_21.4_ = 2.55; P = 0.0184). Moreover, there was no significant decrease to insert fragment size (Figure 3A; T-test; t_32.7_ = 0.952; P = 0.348), genes with mapped sequences (Figure 3C; T-test; t_28.7_ = -0.934; P = 0.358) or total “supercontig” bases recovered (Figure 3D; T-test; t_28.4_ = 0.247; P = 0.807) when libraries were miniaturized from 0.5X to 0.1X reaction volumes.

**Figure 2.**
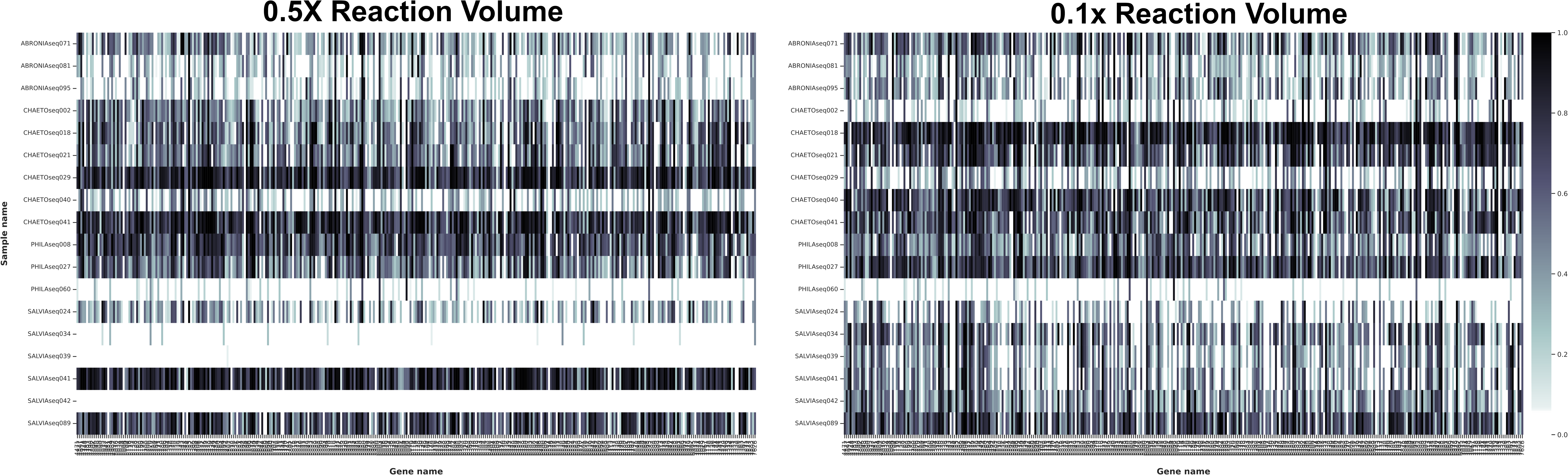
Libraries prepared at O.lX reaction volumes do not have a clear pattern of differentiation in gene recovery when compared to O.SX reaction volumes. The heatmaps show the percent of each gene (column) recovered per sample (row). The left depicts recovery of sequences from the Overlap Set created with 0.5X library preparation reaction volumes, while the right is O. lX, with the Overlap Set samples in the same order top to bottom. The shading of each cell represents the percent gene length recovered for each targeted gene out of 353 total genes. Per the figure legend, the blaclc cells in the heatmap indicate 100% recovery while white cells indicate 0% recovery.

**Figure 3.**
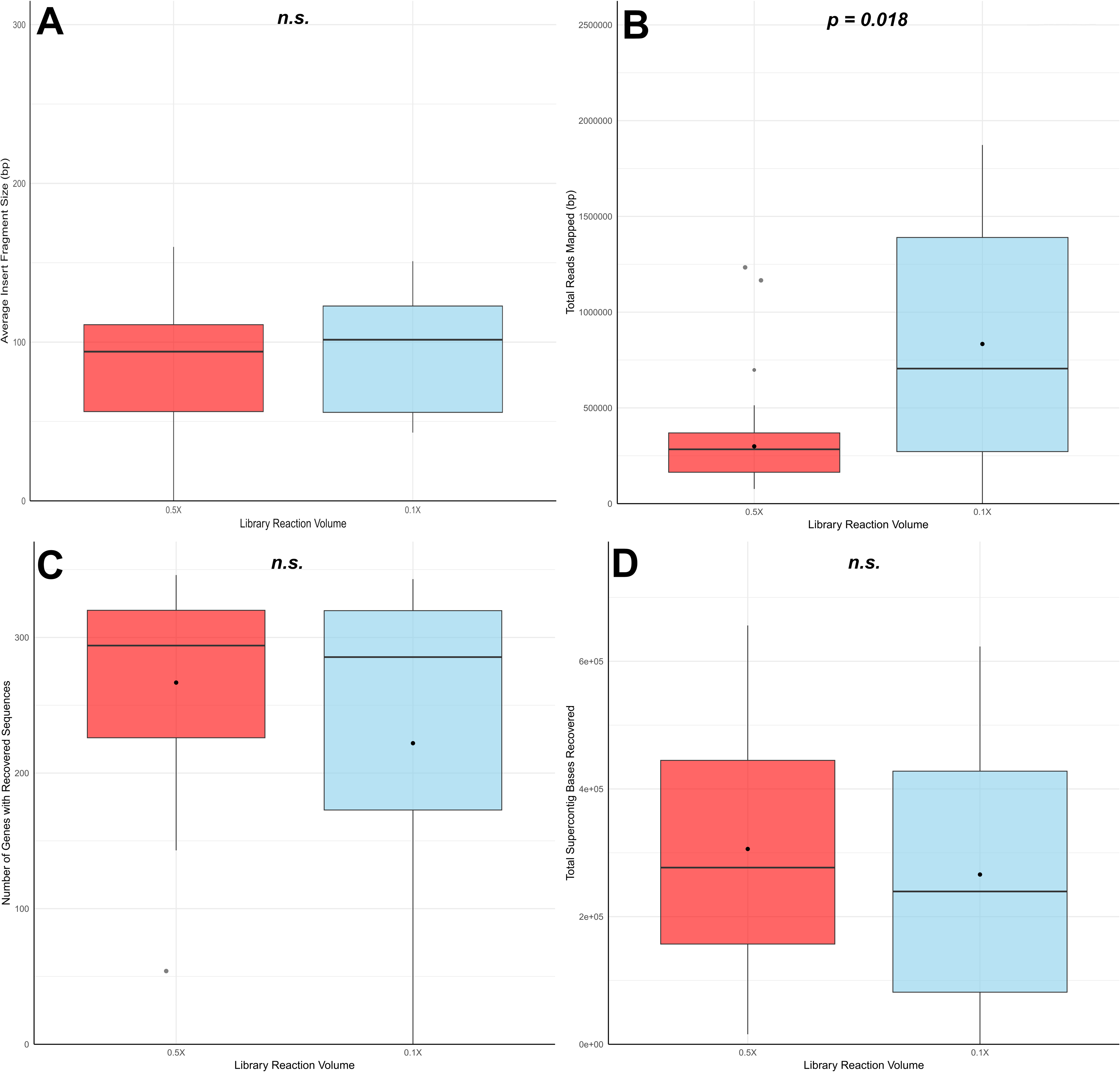
Library miniaturization has no significant negative effects on post-sequencing statistics including reads mapped, genes with recovered sequences, and total number of “supercontig” base pairs recovered in the Overlap Set. A. T-test; t32.7 = 0.952; P = 0.348. B. T-test; t21.4 = 2.55; P = 0.0184. C: T-test; t28.7 = -0.934; P = 0.358. D. T-test; t28.4 = 0.247; P = 0.807.

We also aimed to define any significant differences between miniaturized libraries prepared from fresh tissue-derived DNA and from herbarium tissue-derived DNA using the complete 768 Half-Volume Only and 768 Tenth-Volume Only dataset. Despite finding significant differences in insert fragment sizes (Figure 4A; ANOVA; F_3, 1517_ = 95.43; P < 0.001), total reads mapped (Figure 4B; ANOVA; F_3, 1517_= 2.893; P = 0.034), and genes with mapped sequences (Figure 4C; ANOVA; F_3, 1517_ = 25.0; P < 0.001), direct comparisons via Tukey HSD tests between specific tissue type/reaction volume pairs varied (Supplementary Figure 1). For example, there was no significant difference in insert fragment size for Tenth-Volume Only herbarium and Tenth-Volume Only fresh-tissue derived libraries or genes with sequences between Half-Volume Only fresh-tissue libraries and both Tenth-Volume Only herbarium and fresh tissue libraries. In addition, when comparing reads mapped and genes with sequences for all Half-Volume Only herbarium-derived samples to Tenth-Volume Only herbarium-derived samples, there was no significant difference. These results were largely reflected in the two-way ANOVA as well. However, in terms of reads mapped, the two-way ANOVA attributed differences primarily to tissue type rather than library reaction volume or an interaction between both effects (Supplementary Table 1).These results suggest that library miniaturization is a viable method for target capture, and miniaturized libraries of herbarium tissue-derived DNA perform just as well as half-volume herbarium libraries.

**Figure 4.**
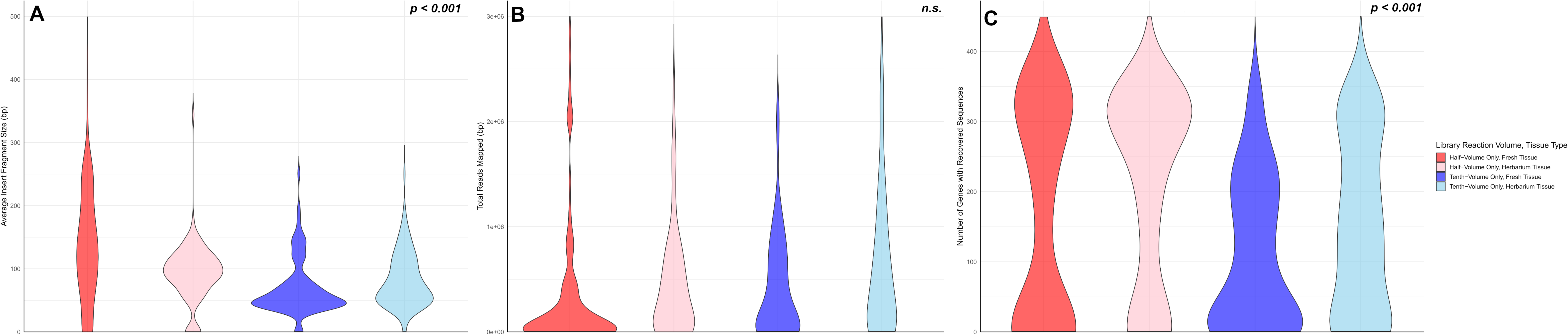
Library miniaturization and tissue type contribute significantly to the variance in average insert fragment size (A. ANOVA; F3, 1517 = 95.43; P < 0.001), total reads mapped (B. ANOVA; F3, 1517= 2.893; P = 0.034) and number of genes with recovered sequences (C. ANOVA; F3, 1517 = 25.0; P < 0.001). However, not all groups are significantly different in each category from each other, based on the Tukey Honestly Significant Differences test (Supplementary Figure 1).

Plastid gene recovery from a subset of the half volume and tenth volume samples, including the Overlap Set libraries was variable and generally low, especially with half-volumes.

The percentage of reads mapping to plastid targets reached a maximum of 2.2%, with 31% of 0.5X samples (Supplementary Figure 4) and 13% of 0.1X samples (Supplementary Figure 5) recovering no plastid genes. The samples with zero gene recovery all had fewer than 5,000 reads mapped to the plastid targets. Above 5,000 reads mapped, gene recovery was more consistent (Supplementary Figure 6), exceeding 40 genes recovered in five 0.1X and 18 0.5X samples. As with nuclear gene recovery, there were several samples where gene recovery was high in one library prep volume but not the other (Supplementary Figures 4&5).

To compare relative costs associated with various components of library preparation across reaction volumes, we calculated the costs of consumables, library preparation reagents, and quality control reagents used in 1.0X, 0.5X, and 0.1X library preparation reactions to the proportion of the cost they represent relative to that cost in 1.0X volumes (Table 1). We concluded that 0.1X library miniaturization decreases costs compared to both 0.5X and 1.0X across all three expense categories, with the largest reduction occurring in library preparation reagents. We also found that utilization of robot automation decreases on-hand temporal costs by more than 4-fold. Therefore, we believe that library miniaturization is an effective method for cost-reduction in both finances and time in target capture sequencing.

**Table 1.** Libraries prepared at O.lX reaction volumes cost less than the libraries prepared at O.SX and I.OX volumes across all components. “Consumables” includes items such as pipette tips, microcentrifuge tubes, and PCR plates. “Library Preparation Reagents” includes the reagents involved in preparing the libraries. “Quality Control Supplies” includes the reagents and consumables used in confirming library concentration and quality. Note that these costs are specific to Texas Tech University and may not be universally applicable. In addition, these totals do not include the costs of DNA extraction or sequencing, only library preparation.

|  | Full Library Reaction Volume |  | Half Library Reaction Volume |  | Tenth Library Reaction Volume |  |
| --- | --- | --- | --- | --- | --- | --- |
| Hands-On Library Prep<br>Time per Plate | ~ 14 hours |  | ~ 14 hours |  | ~ 3 hours |  |
| Item | Per Sample | Per Plate | Per Sample | Per Plate | Per Sample | Per Plate |
| Consumables | \$9.18 | \$881.68 | \$9.18 | \$881.68 | \$2.17 | \$208.67 |
| Library Preparation Reagents | \$26.56 | \$2,550.18 | \$13.28 | \$1,275.09 | \$2.66 | \$255.02 |
| Quality Control Supplies | \$1.00 | \$96.01 | \$0.79 | \$75.72 | \$0.68 | \$65.75 |
| **Totals** | **\$36.75** | **\$3,527.88** | **\$23.26** | **\$2,232.49** | **\$5.51** | **\$529.44** |

## DISCUSSION

We asked whether library miniaturization for targeted sequencing of angiosperm DNA derived from both fresh and herbarium tissue is a valid method for cost reduction. The primary impact of the reduced reaction volumes was a significant decrease in insert sizes. Despite this, when comparing the same set of samples across both reaction volumes, we found that decreased reaction volumes do not significantly reduce the number of genes with recovered sequences, reads mapped, or total length of recovery. Therefore, we determined that library miniaturization is as effective for herbarium tissue-based DNA as it is for fresh tissue-based DNA. In addition, we found that total per-sample cost of 0.1X library miniaturization amounts to less than a quarter of total per-sample costs for 1.0X reactions. We conclude that library miniaturization is a legitimate method for cost reduction of angiosperm targeted sequencing.

To our knowledge, few studies have used library miniaturization on plant tissues for sequencing, and no studies have been conducted utilizing library miniaturization with plant target capture sequencing or herbarium DNA input. Ogiso-Tanaka et al. (2018) had success with 0.1X soybean library miniaturization prior to AmpliSeq sequencing, a reduced representation sequencing method like targeted sequencing. They found that 0.12X was the minimum volume able to produce stable reactions of the correct amplicon. They also concluded that library miniaturization via automated robot liquid handling reduced financial costs by 86.8% and amount of time taken by 85%. However, they only used fresh tissue for their DNA extractions, not antique DNA samples extracted from herbarium specimens like those utilized in our study (Ogiso-Tamaka et al., 2018). Because herbaria are instrumental historical records of global plant distribution and diversity, the ability to cost-effectively use them for genetic analyses opens a new realm of possibilities for botanical research. In addition, our protocol for herbarium tissue resulted in a 38.9% decrease in amount of time to perform library preparation.

Unexpectedly, we found there was no significant difference in reads mapped from Tenth- Volume Only libraries compared to Half-Volume Only samples (Figure 1B). We believe this is likely because insert fragment sizes were shorter on average in the Tenth-Volume Only libraries when compared to the Half-Volume Only libraries. It is possible that because there was no change in the overall number of reads between Tenth-Volume Only and Half-Volume Only libraries, there is a higher number of shorter fragments to account for lack of change. Those shorter fragments have the possibility to overlap with one another more when aligned, leading to higher read depth, and subsequently more reads mapped overall when compared to Half-Volume only libraries. While they do not seem to affect the quality of targeted sequencing, they may negatively impact library preparation for other sequencing methods that require longer reads, such as whole genome sequencing. Future research should focus on exploring the potential effects of shorter insert fragment sizes and ways to mitigate the insert size reduction during miniaturization.

Our study identified variable recovery of plastid loci from both library volumes. The initial description of HybSeq (Weitemeir et al., 2014) and many of the first studies using target capture in plants included genome skimming as part of the advantages of the approach. More recently, Pokorny et al. (2024) reported a median recovery of two-thirds of plastid genes in the Papaveraceae but reported over 1 million reads mapped to their nuclear targets. We were hopeful that the deeper sequencing on the NovaSeq 6000 would match this output, but our results for plastid genes fell short of Pokorny et al. 2024. As advancements in the MyBaits protocol since 2014 have increased target capture efficiency, it potentially has impacted the utility of off-target reads. We would recommend that studies where organellar data is an important component should consider using the “spiking” method to add unenriched libraries to the enriched libraries prior to sequencing, to bias the off target reads for high-copy DNA like organellar genes.

This study did have its limitations, primarily due to using an opportunistic dataset produced from other projects. While there were equal numbers of Half-Volume Only and Tenth- Volume Only libraries sequenced, not all 768 samples of one sequencing run were identical to the 768 samples of the other. In the full dataset, we see a significant reduction in genes recovered in the Tenth-Volume Only libraries; however, this appears to be driven entirely by the Half-Volume Only herbarium samples, which had an unexpectedly high number of genes recovered. In contrast, the 18 individual samples within our Overlap Set were sequenced at both reaction volumes and show no significantly negative effects of the 0.1X library preparation method in terms of insert fragment size, reads mapped, genes with sequences, or total base pairs recovered in the “supercontigs.” So, it is possible that the significant differences between the Half-Volume Only and Tenth-Volume Only datasets are due to stochastic differences in the samples themselves and not sequencing method or tissue type.

Also, due to the opportunistic dataset, we did not have equal numbers of fresh tissue and herbarium tissue-derived DNA samples. Because the samples in our Overlap were only derived from herbarium tissue, we addressed this with a power analysis via comparison of 100 random herbarium samples to 100 random fresh tissue samples for both 0.5X and 0.1X reaction volumes, amounting to 400 samples total. The power analysis results largely mirrored our total dataset results. For example, Half-Volume Only fresh tissues samples had significantly higher insert sizes than herbarium samples at both reaction volumes. This is to be expected, as herbarium tissue is often already degraded due to age. However, our power analysis results did differ in a few key ways: comparisons of insert fragment size between Half-Volume Only and Tenth- Volume Only herbarium libraries, reads mapped in Half-Volume Only fresh and Tenth-Volume Only herbarium libraries, and genes with sequences in Half-Volume fresh and herbarium were found non-significant via one-way ANOVA and subsequent post-hoc Tukey HSD testing (Supplementary Figures 2-3). These differences between the full dataset and this power analysis subset lead us to believe that sampling balance and the opportunistic nature of the data set influences significance patterns in the tissue type and library reaction volume interactions, and the biological trend is more consistent in the balanced subset: 0.1X fresh loses some performance, herbarium DNA is volume-sensitive but not as dramatically impacted.

Although the balanced subset analysis revealed that 0.1X fresh libraries recovered significantly fewer genes with sequences than 0.5X libraries, performance losses were modest and not consistently observed across herbarium samples. Importantly, herbarium libraries prepared at 0.1X volumes performed comparably to 0.5X fresh libraries, suggesting that degradation status of input DNA has a stronger influence than reaction volume. Taken together, these results demonstrate that while some contexts may benefit from half-volume reactions, miniaturization to 0.1X remains a broadly valid strategy for substantial cost reduction with only minor impacts on data recovery. Finally, our study focused on the Angiosperms353 target capture probe set. For the broadest utilization of our protocol, we would recommend future research confirm the effectiveness of library miniaturization for taxon-specific probe sets such as Compositae1061 (Asteraceae; Mandel et al., 2014), Cactaceae591 (Cactaceae; Romeiro-Brito et al., 2022), GoFlag451 (flagellate land plants; Breinholt et al., 2021), and Nikolov1827 (Brassicaceae; Nikolov et al., 2019).

In summary, we determined the efficiency of library miniaturization for cost reduction of both angiosperm DNA sequencing from both fresh and herbarium tissue-derived via the comparison of two reduced reaction volumes: 0.5X and 0.1X. We found that post-sequencing data quality generated by both methods as well as both tissue types differed via decreased insert fragment size, decreased genes with recovered sequences, and increased reads mapped in 0.1X reactions. These results support our hypothesis that library miniaturization can be used for cost reduction of both fresh and herbarium sample targeted sequencing without major reduction in post-sequencing data quality. We hope this study further shows the importance and value of utilizing herbarium and general natural history collection data in conservation and other multi-taxa studies as a cost reduction method. By decreasing the costs of genetic techniques, we increase the accessibility of sequencing population-level datasets to groups with lower funding sources. This could greatly benefit future conservation management plans where integration of genetic data is otherwise infeasible.

## Acknowledgements

We thank Sherese Price and Madeline Slimp for providing pre-publication access to samples, and Haley Hale for valuable advice on laboratory protocols. We are grateful to Tigga Kingston, Ari Rice, Anastasia Chouvalova, Victor Kayejo, and Sean Sutor for their careful review of the manuscript. We also acknowledge the personnel of SPTLabtech for their technical support. Finally, we thank the anonymous reviewers for their insightful, constructive feedback. This research was supported by the Center for Biotechnology and Genomics at Texas Tech University.

## Author Contributions

MRB and MGJ designed the study and developed methodology with input from MF. MF also secured funding for the equipment utilized in this project. MRB performed the wet laboratory components as well as collected and organized the data. MRB, RDC, and EAS analyzed the data and prepared figures and tables. MRB wrote the original draft of the manuscript. All authors contributed to manuscript review and approved the final submission.

## Data Availability

Sequencing statistics spreadsheets for the Half-Volume, Tenth-Volume, and Overlap data sets as well as the R Code utilized in the analysis are available in a Dryad repository at https://doi.org/10.5061/dryad.4mw6m90qf. In addition, we will upload sequence files for the Overlap Set to the Sequence Read Archive (SRA) upon acceptance of this manuscript.

