## Supplementary Table 1 for "Fun-Sized Library Prep: Miniaturization is a valid method for per-sample cost reduction in targeted sequencing of angiosperm DNA"

***Supplementary Table 1.*** Two-way ANOVA table comparing the main effects (tissue type and library reaction volume) as well as their interaction for significance.

| | | Degrees of Freedom | Sum of Squares | Mean Square | F-statistic | P-value |
|---|---|---|---|---|---|---|
| **Average Insert Fragment Size (bp)** | Tissue Type | 1 | 1000995 | 1000995 | 34.55 | <0.001 |
| | Library Reaction Volume | 1 | 491371 | 491371 | 168.08 | <0.001 |
| | Tissue Type : Library Reaction Volume | 1 | 242730 | 242730 | 83.03 | <0.001 |
| | Residuals | 1517 | 4434796 | 2923 | | |
| **Total Reads Mapped** | Tissue Type | 1 | 9.55E+12 | 9.55E+12 | 6.916 | 0.009 |
| | Library Reaction Volume | 1 | 3.46E+12 | 3.46E+12 | 2.504 | 0.114 |
| | Tissue Type : Library Reaction Volume | 1 | 2.68E+11 | 2.68E+11 | 0.194 | 0.660 |
| | Residuals | 1517 | 2.10E+15 | 1.38E+12 | | |
| **Number of Genes with Recovered Sequences** | Tissue Type | 1 | 371678 | 371678 | 22.634 | <0.001 |
| | Library Reaction Volume | 1 | 772890 | 772890 | 47.066 | <0.001 |
| | Tissue Type:Library Reaction Volume | 1 | 100878 | 100878 | 6.143 | 0.013 |
| | Residuals | 1517 | 24911186 | 16421 | | |
