## Supplementary Table 2 for "Fun-Sized Library Prep: Miniaturization is a valid method for per-sample cost reduction in targeted sequencing of angiosperm DNA"

***Supplementary Table 2.*** Voucher information for herbarium specimens in the Overlap Set sampled for this study. Specimens are housed in the University of Arizona Herbarium (ARIZ), Arizona State University Vascular Plant Herbarium (ASU), Desert Botanical Garden Herbarium (DES), New Mexico State University Herbarium (NMC), Texas Tech University E. L. Reed Herbarium (TTC), University of New Mexico Herbarium (UNM), and University of Texas at El Paso Biodiversity Collections Herbarium (UTEP).

| Herbarium Barcode | Herbarium | Scientific Name | Collector | Collection Numbe | Collection Date |
|---|---|---|---|---|---|
| UNM0059436 | University of New Mexico Herbarium | *Abronia fragrans* | F. Broeske | FO-9-A | 4/24/1965 |
| ASU0114382 | Arizona State University Vascular Plant Herbarium | *Abronia fragrans* | R. L. Tresler | 359 | 7/10/1967 |
| 38701 | New Mexico State University Herbarium | *Abronia fragrans* | R. W. Allen | 2 | 5/8/1949 |
| TTC009677 | Texas Tech University, E. L. Reed Herbarium | *Chaetopappa ericoides* | Demaree | 7683 | 5/14/1930 |
| SJNM-V-0008857 | San Juan College Herbarium | *Chaetopappa ericoides* | Goodrich, S. | 23704 | 5/7/1992 |
| ASU0296124 | Arizona State University Vascular Plant Herbarium | *Chaetopappa ericoides* | J. André | 22449 | 5/24/2012 |
| TTC009672 | Texas Tech University, E. L. Reed Herbarium | *Chaetopappa ericoides* | Tommy Rosson | 1357 | 5/11/1968 |
| DES00084090 | Desert Botanical Garden Herbarium | *Abronia fragrans* | Jim André | 23715 | 4/26/2012 |
| 292572 | University of Arizona Herbarium | *Chaetopappa ericoides* | P.S. Martin | s.n. | 6/7/1970 |
| UTEP:Herb:16536 | University of Texas at El Paso Biodiversity Collections H | *Philadelphus mearnsii* | Thomas R. Van Devender | - | 6/16/1981 |
| 50437 | New Mexico State University Herbarium | *Philadelphus mearnsii* | R. W. Spellenberg, D. E. Ward | 5522 | 5/13/1980 |
| TTC021780 | Texas Tech University, E. L. Reed Herbarium | *Philadelphus hitchcockian* | T.L. Burgess | 1142 | 7/7/1973 |
| UTEP:Herb:16197 | University of Texas at El Paso Biodiversity Collections H | *Salvia summa* | Thomas R. Van Devender, Christine Le | - | 6/23/1981 |
| UTEP:Herb:16185 | University of Texas at El Paso Biodiversity Collections H | *Salvia farinacea* | Carl S. Lieb | Lieb 137 | 7/4/1981 |
| TTC014140 | Texas Tech University, E. L. Reed Herbarium | *Salvia farinacea* | B.G. Cumbie | 180 | 5/29/1961 |
| TTC014151 | Texas Tech University, E. L. Reed Herbarium | *Salvia farinacea* | Dana, S.W. | 25 | 5/24/1976 |
| ASU0111349 | Arizona State University Vascular Plant Herbarium | *Salvia azurea* | L.C. Higgins | 6264 | 9/2/1972 |
| TTC020517 | Texas Tech University, E. L. Reed Herbarium | *Salvia farinacea* | D. K. Northington | 528 | 6/6/1973 |
