## Supplementary Figure 1 for "Fun-Sized Library Prep: Miniaturization is a valid method for per-sample cost reduction in targeted sequencing of angiosperm DNA"

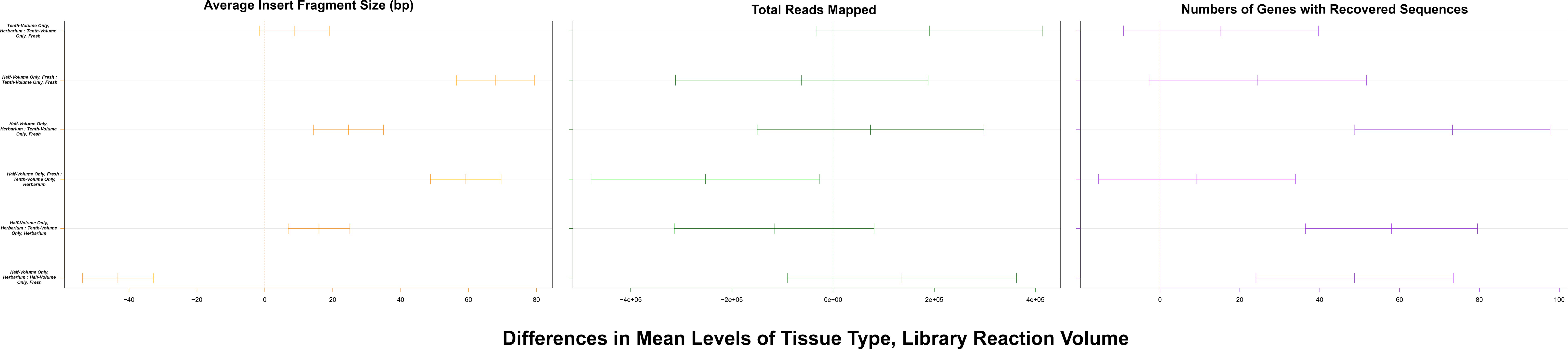


***Supplementary Figure 1.*** Tukey Honestly Significant Difference plots depicting which tissue type/library reaction volume pairs using the Half-Volume Only and Tenth-Volume Only datasets are significantly different from one another. The whiskers of each plotted line refer to the 95% confidence intervals. If the line crosses over the zero mark, the groups are not significantly different from each other. If the whiskers do not cross over the zero mark and the entire line is less than zero, then the first group is significantly lower than the second group. However, if the entire line is greater than zero, then the first group is significantly higher than the second group.
