## Supplementary Figure 2 for "Fun-Sized Library Prep: Miniaturization is a valid method for per-sample cost reduction in targeted sequencing of angiosperm DNA"

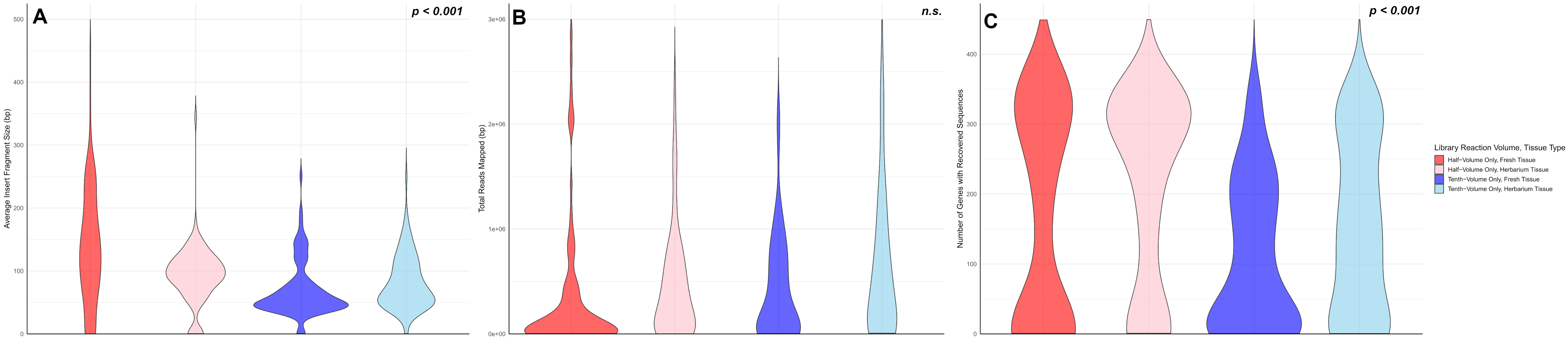


***Supplementary Figure 2.*** Power Analysis of a subset of 100 samples from each library reaction volume-tissue type combination provides similar results as total Half-Volume Only and Tenth-Volume Only datasets. Metrics utilized are average insert fragment size in base pairs (A. ANOVA; F-value = 22.84; $P < 0.001$; df = 3), total reads mapped (B. ANOVA; F-statistic = 0.602; $P = 0.602$; df = 3) and number of genes with recovered sequences (C. ANOVA; F-statistic = 9.793; $P < 0.001$; df = 3) between library reaction volume/tissue type groups. However, not all groups are significantly different in each category from each other, based on the Tukey Honestly Significant Differences test (Supplementary Figure 3).
