## Supplementary Figures 4-5 for "Fun-Sized Library Prep: Miniaturization is a valid method for per-sample cost reduction in targeted sequencing of angiosperm DNA"

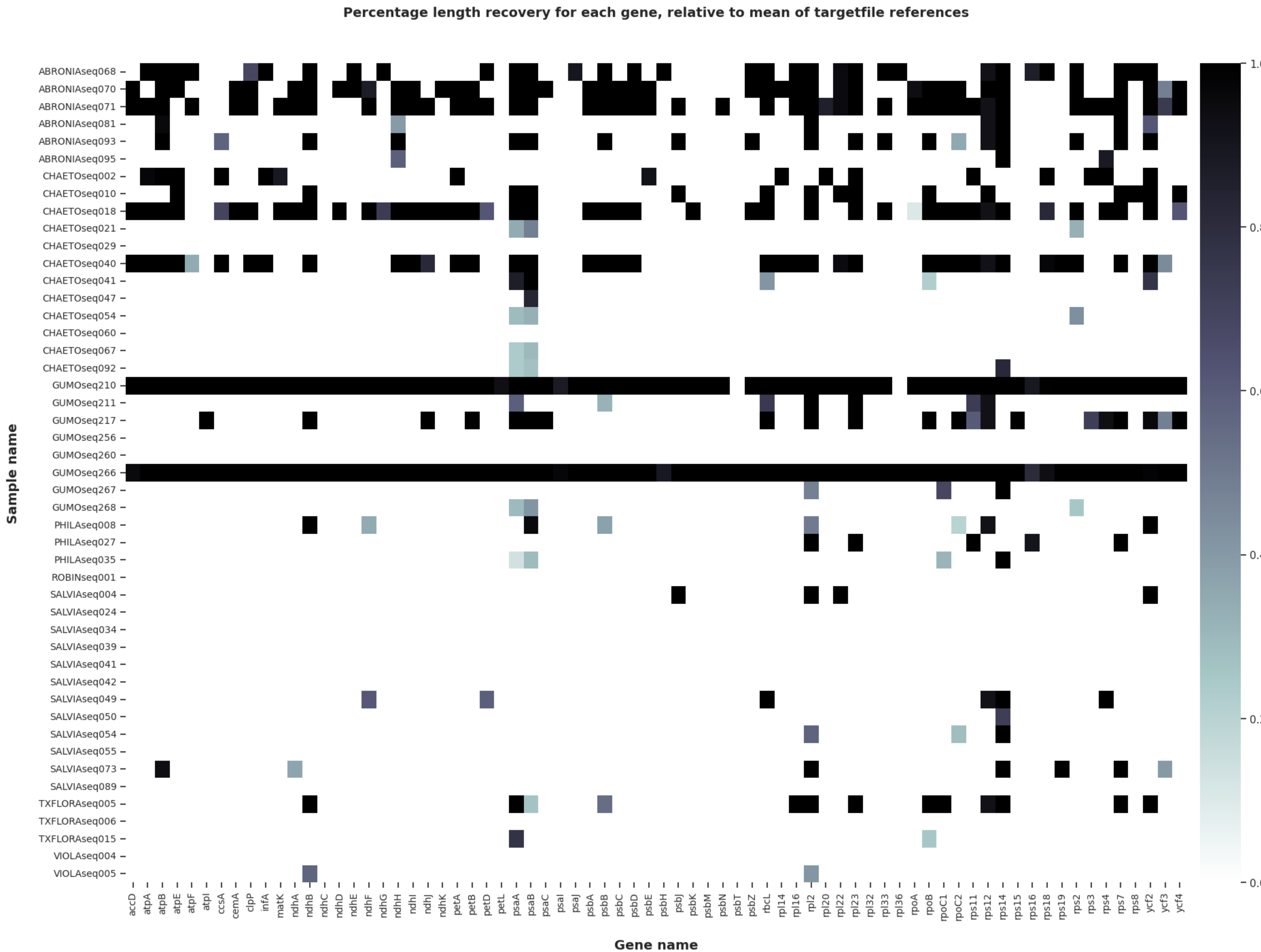


***Supplementary Figure 4.*** Heatmap from HybPiper shows recovery of plastid genes from the off-target reads generated through targeted sequencing of Angiosperms353 with half-volume library preparation. Each row is a sample and each column is a gene. The shading of each cell is the percentage of the expected length of the gene recovered

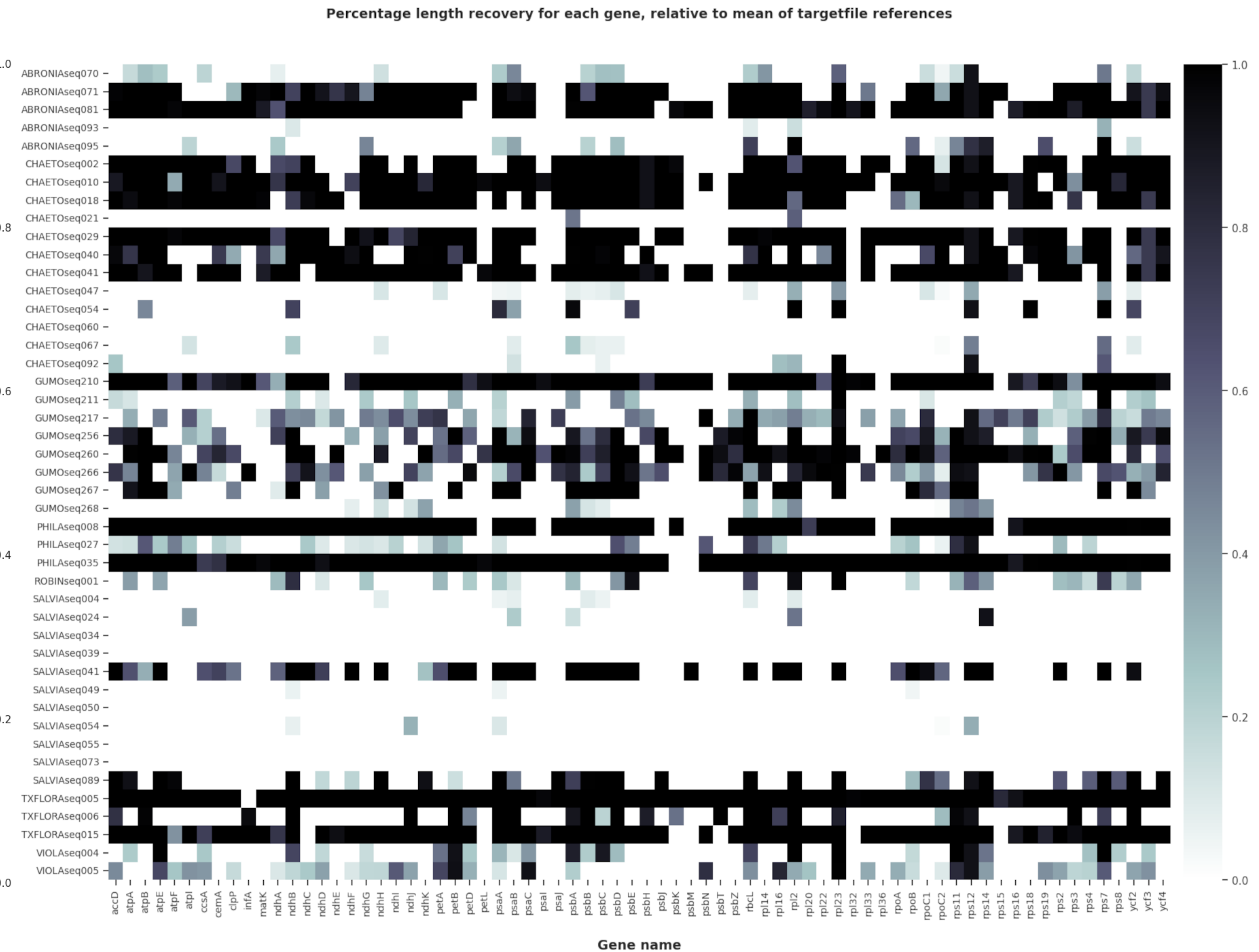


***Supplementary Figure 5.*** Heatmap from HybPiper shows recovery of plastid genes from the off-target reads generated through targeted sequencing of Angiosperms353 with tenth-volume library preparation. Each row is a sample and each column is a gene. The shading of each cell is the percentage of the expected length of the gene recovered.
