## Supplementary Figure 6 for "Fun-Sized Library Prep: Miniaturization is a valid method for per-sample cost reduction in targeted sequencing of angiosperm DNA"

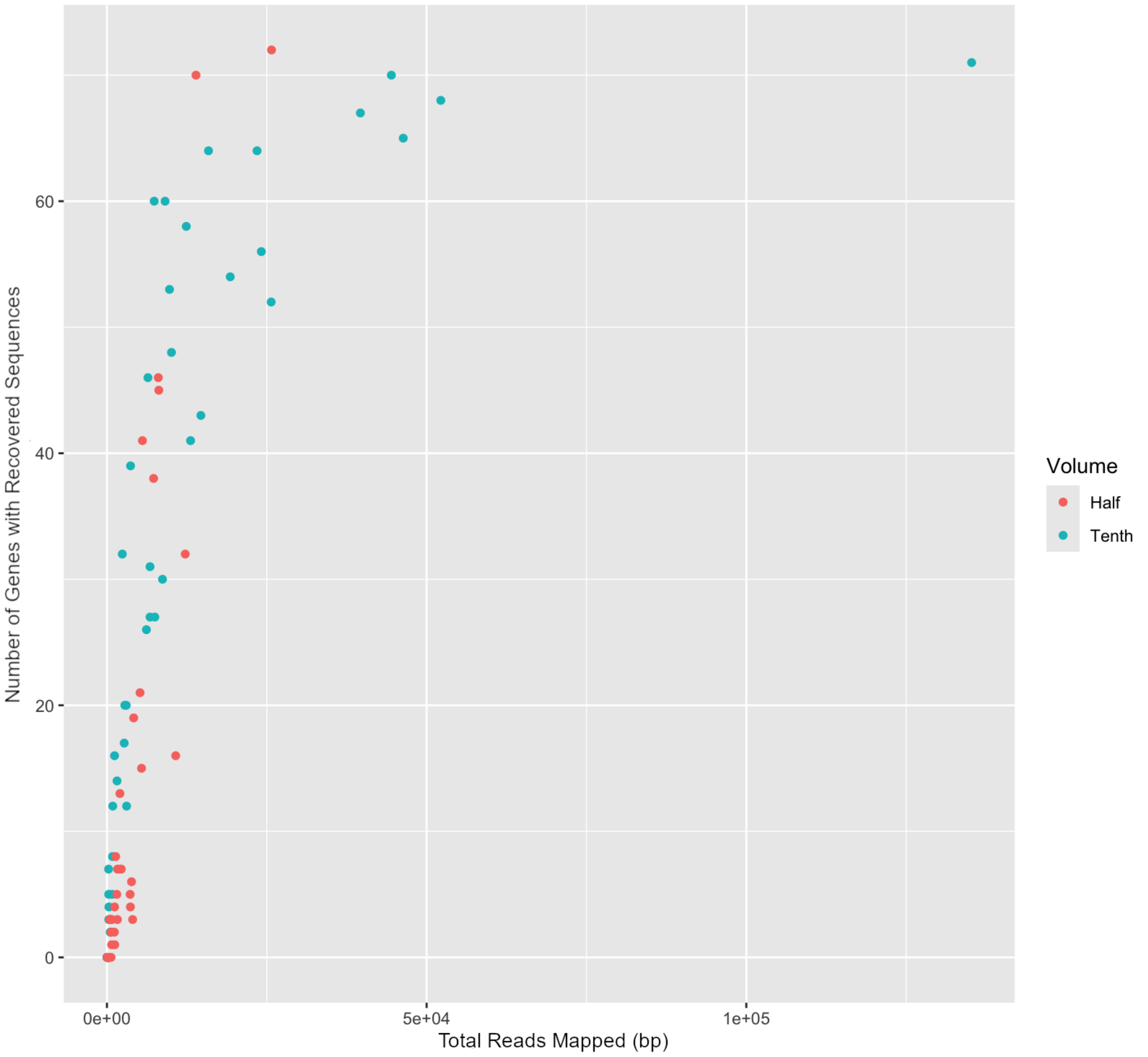


***Supplementary Figure 6.*** Scatterplot showing the relationship between the number of reads mapped to plastid genes and the number of plastid genes with sequences, using either half-volume (red) or tenth-volume (blue) library preparation.
